# Spatiotemporal patterns of high-frequency activity (80-170 Hz) in long-term intracranial EEG

**DOI:** 10.1101/2020.03.26.999425

**Authors:** Zhuying Chen, David B. Grayden, Anthony N. Burkitt, Udaya Seneviratne, Wendyl J. D’Souza, Chris French, Philippa J. Karoly, Katrina Dell, Mark J. Cook, Matias I. Maturana

**Author notes:** Corresponding author: Ms. Zhuying Chen, Department of Biomedical Engineering, The University of Melbourne. Address: 203 Bouverie Street, Melbourne, VIC, Australia, 3053. There are supplementary tables and figures for the manuscript. Search terms: [62] EEG, [65] Intracranial electrodes, [70] Epileptogenic zone, [77] Partial seizures, [81] Complex partial seizures.

## Abstract

**Objective:** To assess the variability in the rates and locations of high-frequency activity (HFA) and epileptiform spikes after electrode implantation, and to examine the long-term patterns of HFA using ambulatory intracranial EEG (iEEG) recordings.

**Methods:** Continuous iEEG recordings obtained over an average of 1.4 years from 15 patients with drug-resistant focal epilepsy were used in this study. HFA was defined as high-frequency events with amplitudes clearly larger than the background, which was automatically detected using a custom algorithm. High-frequency oscillations (HFOs) were also visually annotated by three neurologists in randomly sampled segments of the total data. The automatically detected HFA was compared with the visually marked HFOs. The variations of HFA rates were compared with spikes and seizures on patient-specific and electrode-specific bases.

**Results:** HFA was a more general event that encompassed HFOs manually annotated by different reviewers. HFA and spike rates had high amounts of intra- and inter-patient variability. The rates and locations of HFA and spikes took up to weeks to stabilize after electrode implantation in some patients. Both HFA and spike rates showed strong circadian rhythms in all patients and some also showed multiday cycles. Furthermore, the circadian patterns of HFA and spike rates had patient-specific correlations with seizures, which tended to vary across electrodes.

**Conclusions:** Analysis of HFA and epileptiform spikes should account for post-implantation variability. Like seizures, HFA and epileptiform spikes show circadian rhythms. However, the circadian profiles can vary spatially within patients and their correlations to seizures are patient-specific.

## Introduction

Over the past two decades, high-frequency activity in electroencephalography (EEG) with frequencies above 80 Hz has received significant attention in epilepsy research. This activity appears to increase in intensity before and during seizures and these increases are predominantly localized in the seizure onset zone (SOZ).^1–4^ A subtype of high-frequency activity (HFA), termed high-frequency oscillations (HFOs), has been proposed as a biomarker of the epileptogenic zone (EZ).^5^ HFOs occur more frequently in the SOZ,^6–8^ and removing brain areas exhibiting HFOs is correlated with a good surgical outcome.^9–11^

However, the literature shows contradictory results regarding HFOs.^8, 12^ This may be partly due to post-implantation effects, which cause changes in intracranial EEG (iEEG) signals lasting months.^13, 14^ No studies have yet determined whether EEG biomarkers show similar variations, which is an important subject as clinical decisions based on biomarkers extracted immediately following implantation could be inaccurate.

Manual review of EEG is the “gold standard” for HFO identification, but is only feasible with small datasets and has poor agreement among reviewers.^15–17^ Given the lack of a clear definition of HFOs^17^ and the impracticality of performing manual review of HFOs in long-term data, we devised a more general definition of high-frequency activity that encompasses HFOs and other high-frequency events that are typically detected by automatic techniques.^16, 18–20^

This study shows that the behavior of HFA and epileptiform spikes immediately after implantation of electrodes does not reflect the long-term behavior after recovery from surgery. We also show that HFA rates fluctuate with periodicities of similar duration to seizures.

## Methods

### Data source

This study used long-term iEEG recordings from 15 patients with drug-resistant focal epilepsy participating in a clinical trial of a seizure advisory system.^21^ Patient demographics and recording durations can be seen in Table 1. The recording device comprised 16 electrodes implanted over the presumed EZ based on prior lesion imaging or seizure history. Data were sampled at 400 Hz. In total, 3273 seizures (average 204 seizures per patient, range 9-545 seizures) were recorded over an average of 519 days (range 184-767 days). Seizure start and end times were annotated by expert neurologists. Seizures with clinical symptoms (type 1 seizures) and electrographic seizures identical to type 1 seizures but without clinical symptoms (type 2 seizures) were included in this study. Epileptiform spikes (herein termed spikes) were also analyzed in this study; the detection of spikes has been previously validated.^22^

**Table 1.** Patient characteristics at the baseline and the detected rates of HFA and spikes

| Pt No. | Age (year) | Gender | Duration (day) | HGE rate (/min) |  | Spike rate (/min) |  | seizure counts | EZ |
| --- | --- | --- | --- | --- | --- | --- | --- | --- | --- |
|  |  |  |  | Mean* | CV* | Mean* | CV* |  |  |
| 1 | 26 | M | 767 | 3.32 [2.27,11.79] | 1.29 [1.08,1.54] | 0.10 [0.02,3.15] | 5.37 [1.63,13.64] | 151 | PT |
| 2 | 44 | M | 730 | 1.34 [0.17,32.69] | 1.93 [0.96,2.73] | 0.09 [0.00,0.84] | 2.97 [1.37,13.64] | 32 | OP |
| 3 | 22 | F | 557 | 20.37 [5.46,41.69] | 0.84 [0.68,1.16] | 0.71 [0.00,3.51] | 2.94 [1.10,18.81] | 390 | PT |
| 4 | 61 | M | 232 | 0.59 [0.05,5.29] | 2.71 [0.87,6.21] | 0.68 [0.01,36.49] | 1.55 [0.67,4.96] | 22 | PT |
| 5 | 20 | F | 272 | 0.88 [0.07,8.25] | 1.72 [0.75,3.99] | 0.12 [0.00,1.79] | 2.08 [0.87,6.24] | 9 | FT |
| 6 | 62 | M | 441 | 5.04 [0.17,13.23] | 1.61 [1.12,2.72] | 0.93 [0.01,22.60] | 1.30 [0.80,6.15] | 71 | T |
| 7 | 52 | M | 184 | 0.79 [0.22,10.32] | 1.83 [0.81,2.64] | 1.88 [0.05,22.22] | 1.27 [0.58,3.10] | 313 | FT |
| 8 | 48 | M | 558 | 3.67 [1.84,44.70] | 1.50 [0.38,1.92] | 3.41 [0.29,24.80] | 0.90 [0.39,1.69] | 467 | FT |
| 9 | 51 | F | 394 | 1.79 [0.35,30.46] | 1.45 [0.63,2.99] | 1.07 [0.05,5.56] | 2.37 [0.78,15.31] | 204 | OP |
| 10 | 50 | F | 373 | 6.67 [3.84, 18.72] | 1.34 [0.99, 1.56] | 4.57 [2.30, 8.93] | 1.15 [0.77, 1.82] | 545 | FT |
| 11 | 53 | F | 721 | 2.10 [0.10,8.47] | 1.33 [0.65,2.58] | 1.17 [0.08,16.50] | 1.28 [0.66,22.88] | 467 | FT |
| 12 | 43 | M | 728 | 1.26 [0.12,12.37] | 2.58 [0.75,16.51] | 0.22 [0.01,16.20] | 1.91 [0.62,5.42] | 13 | T |
| 13 | 50 | M | 746 | 7.11 [1.02, 54.81] | 0.87 [0.60, 1.23] | 6.24 [0.26, 26.39] | 0.93 [0.48, 1.16] | 500 | T |
| 14 | 49 | F | 627 | 1.99 [0.56, 12.92] | 1.96 [1.20, 3.64] | 6.68 [0.27, 50.62] | 1.85 [0.79, 6.88] | 12 | PT |
| 15 | 36 | M | 465 | 3.74 [2.33,8.10] | 1.27 [0.85,1.78] | 7.49 [0.11,59.87] | 0.71 [0.43,1.97] | 77 | T |

Pt No = patient number, M = male, F = female, CV = coefficient of variation, EZ = epileptogenic zone, PT = parietal-temporal, OP = occipital-parietal, FT = frontal-temporal, T = temporal. \*Mean and CV are presented as median [minimum, maximum] to show how they were distributed over 16 different electrodes within patients.

### Automated high-frequency activity (HFA) detection and validation

Figure 1 shows a graphical representation of the HFA detection process. The HFA detection process was performed at patient-specific and electrode-specific bases. To detect HFA, iEEG data on each individual electrode was first filtered using an order-12 infinite impulse response (IIR) bandpass filter with cutoff frequencies between 80 and 170 Hz (Figure 1A & 1B). Then, 300 segments each with a duration of 10 minutes were randomly selected from the filtered iEEG. A Gaussian curve was then fit to the histogram of amplitudes of the sampled filtered iEEG (Figure 1C). A threshold of 5 standard deviations (SDs) of the fitted Gaussian curve was selected to detect HFA. The objective was to identify the amplitudes of HFA that substantially differed from the background. The entire time series of filtered iEEG was passed through a Hilbert transform to extract the signal envelope. HFA was detected when the signal envelope crossed the selected threshold (Figure 1D). For each detected event, the time of the event peak, event amplitude, and envelope width were recorded. Events that were less than 100 ms apart were considered to be a single event.

**Figure 1.**
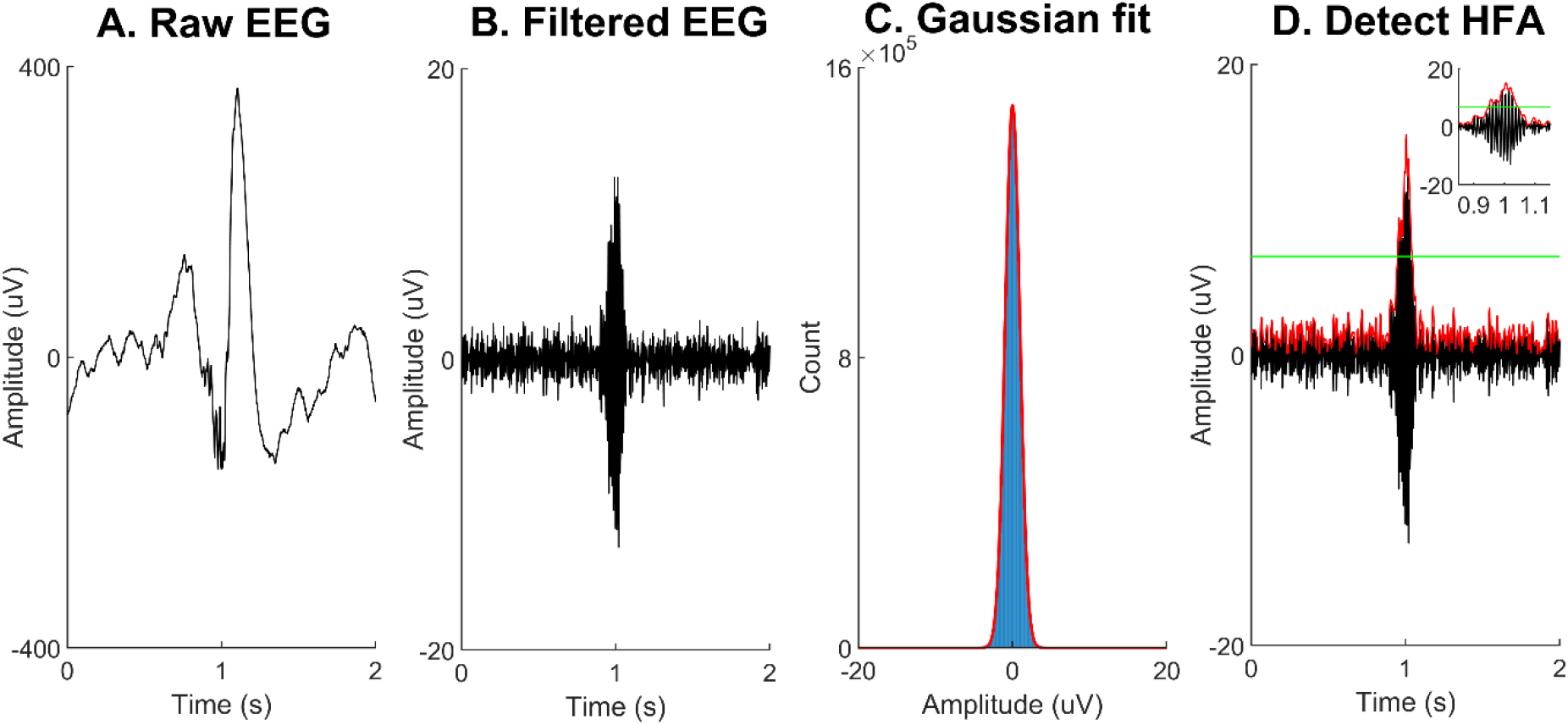
HFA detection procedure. (**A**) Raw iEEG was (**B**) filtered using an order-12 IIR bandpass filter with frequency range 80-170 Hz. (**C**) A Gaussian curve was fitted to the histogram of the amplitudes from 300 randomly selected filtered iEEG samples each of 10 minutes duration. The Gaussian curve was fitted separately for each electrode. (**D**) A threshold of 5 SD of the fitted Gaussian curve was selected as a threshold (green line) to detect HFA. An HFA was detected if the envelope (red curve) exceeded the amplitude threshold.

The dataset contained segments of missing data due to communication dropouts or periods when the device memory was not regularly retrieved. To deal with dropouts, we computed the rates of HFA, which were normalized by the amount of time without dropouts.

To determine the relationship between the detected HFA and manually identified HFOs, automatically detected HFA were compared with HFOs identified by three expert clinical neurologists who were blinded to the selected iEEG segments for HFO marking. For each patient and each recording electrode, we randomly selected 24 samples of iEEG data for HFO reviewing. Each iEEG sample comprised a 1-min epoch, sampled from a randomly selected day such that samples spanned the full 24 hours of a day. For each patient, this produced a total of 384 minutes of iEEG data for independent reviewing. HFO marking was based on the most commonly used criteria of having at least four consecutive oscillations and clearly standing out from baseline.^6, 12, 23^ Markings were performed in a customized MATLAB (MathWorks, R2018b, USA) graphical user interface that we developed, which allowed reviewers to adjust the gain and time window. A total of 360 1-min 16-channels epochs (i.e., 5760 minutes) of iEEG data were randomly shown to each reviewer and reviewers were instructed to mark as many epochs as possible without a time limit. Of this, 112 minutes of iEEG data were reviewed by all reviewers, which were used to measure the interrater agreement.

To evaluate the extent of similarity between automatically detected HFA and visually marked HFOs, we calculated the average proportion of shared events between them, which was computed as

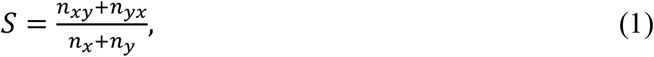

where *S* represents the similarity index, *x* represents HFOs, *y* represents HFA, *n*_*x*_ is the number of HFOs, *n*_*y*_ is the number of HFA, *n*_*xy*_ is the number of HFOs that were also detected as HFA, and *n*_*yx*_ is the number of HFA that were also marked as HFOs. *S* ranges from 0 to 1, where a value close to 1 indicates high similarity and a value close 0 implies low similarity.

The interrater agreement in HFO marking was also computed using the equation (1), where *x* represents HFOs marked by reviewer 1, *y* represents HFOs marked by reviewer 2, *n*_*x*_ is the number of HFOs marked by reviewer 1, *n*_*y*_ is the number of HFOs marked by reviewer 2, *n*_*xy*_ is the number of HFOs marked by reviewer 1 that were also marked by reviewer 2, and *n*_*yx*_ is the number of HFOs marked by reviewer 2 that were also marked by reviewer 1.

### Statistical analysis

Unless stated otherwise, all analyses were conducted using MATLAB. The HFA and spike rates were calculated as average counts per minute per electrode in each 1-hour EEG epoch. For each patient and each electrode, the coefficient of variation was computed to measure the variability of HFA and spike rates in the long-term recordings. The coefficient of variation (CV) was defined as the ratio of the standard deviation (SD) to the mean.

To examine cyclic patterns of HFA and spike rates, the autocorrelation of the rate data was calculated. The autocorrelation function (ACF) measures the similarity of the data with a time-delayed copy of the data as a function of the lag time. The ACF of periodic data shows cyclic patterns with the same period as the data.

To test for the significance of the ACF value at lag *k* (termed *r*_*k*_), we assumed that the signal was a moving average process of order *k* − 1, which means the ACF is theoretically zero at lag ≥ *k*. Then, the standard error (*E*) of *r*_*k*_ can be computed using Bartlett’s formula,^24^

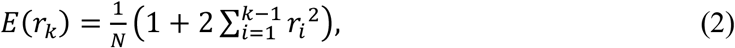

where *N* is the number of samples in the signal. The 99% confidence limits are 0 ± 2.58 × *E*(*r*_*k*_). If *r*_*k*_ is beyond the confidence limits, *r*_*k*_ is statistically unlikely to be caused by chance (*p*<0.01).

To detect the dominant periods in the rate data, an automatic detection method was developed based on the regularity of peaks in the ACF. Since the rate data was often composed of multiple cycles, the rate data was separated into short and long cycles. The separation into short and long cycles and detection of dominant periods were computed in four steps:

1. Filtering: To detect superimposed cycles with different periods within a signal, the rate signal was filtered into short cycles and long cycles using moving average filters. The signal with short cycles was obtained by subtracting the signal smoothed with a moving average filter of length 1 day from the unfiltered signal. The signal with long cycles was obtained by subtracting the signal smoothed with a moving average filter of length 7 days from the signal smoothed with a moving average filter of length 1 day. We considered cycles with durations up to one week.
2. Autocorrelation: The autocorrelation functions (ACFs) of both the short and long cycle signals were computed.
3. Regularity of peaks: Peaks in the ACF of each signal were detected and the mean, SD and CV of the inter-peak intervals were calculated. If CV was less than 0.25, the peaks were considered to be regular, and the mean of inter-peak intervals was selected as the candidate period. Note, to ensure that the period of oscillation was robustly detected, at least 5 prominent peaks in the ACF with similar inter-peak intervals were required.
4. Validation: The candidate period was validated by checking if the ACF value at the nearest integer of the candidate period was outside the 99% confidence limits (equation (2)).

To measure the association of circadian patterns of rates of HFA and spike with seizures, the Spearman correlation coefficient (*R*) was calculated for each patient and each electrode. Nested analysis of variance (ANOVA) was performed using Minitab software (version 19.1) to test whether there were statistically significant variations in correlation values among patients and among electrodes within patients.

### Standard protocol approvals, registrations, and patient consents

Ethics approval for the study was given by the Human Research Ethics Committees of Austin Health, the Royal Melbourne Hospital, and St Vincent’s Hospital in Melbourne, Australia. All patients gave a written informed consent before participation.

### Data availability

A subset of the data used here can be found in the epilepsy ecosystem (www.epilepsyecosystem.org). Data not included in the epilepsy ecosystem can be made available upon reasonable request and subject to ethical approval.

## Results

### HFA and spike rates

For each patient, the mean rates and CVs of the HFA and spike rates were calculated for each of the 16 electrodes (table 1). Overall, the mean rates of HFA and spike were highly variable across patients. For instance, the HFA rate tended to be low for Patient 4 but high for Patient 3, and the spike rate tended to be low for Patient 2 but high for Patient 15. For all patients, the mean rates of HFA and spike exhibited high spatial variability across the 16 electrodes of individual patients. For example, for Patient 13, the electrode with the highest HFA rate had a mean rate of 54 times greater than the mean rate of the lowest rate electrode (54.8 vs. 1.02 /min). For Patient 3, the highest spike rate electrode had a mean rate of 17.6 times greater than the mean rate of the lowest rate electrode (24.1 vs. 1.37/min). Additionally, the HFA and spike rates demonstrated a high amount of temporal variability, evidenced by the consistently high CVs in table 1.

Cross-validation of HFA with HFOs revealed that, among 5760-min epochs, 2688-min epochs were reviewed by by one expert neurologist (US), and 14,666 HFOs were manually marked, while 21,917 HFA events were detected by our detector in these reviewed epochs (table e-1). Pooling all events together, 71% of HFA and HFOs coincided. Among 112-min epochs, which were independently marked by three expert neurologists (US, WD, and CF), the average similarity of HFA and HFOs was 67%, while the average inter-rater agreement in HFO marking was 69% (table e-2).

### Post-implantation variability

Until now, the majority of HFO publications have been based on the analysis of the short (1, 5 or 10-min) sleep iEEG recordings.^12, 19, 23, 25, 26^ However, one recent study has challenged this choice by showing that different 10-min segments of sleep iEEG data produced highly variable HFO rates and locations.^8^ Our previous study also demonstrated that iEEG signals fluctuated over months post-implantation before stabilizing.^13^ Motivated by these results, we examined the evolution of HFA and spike rates post-implantation. An example of the post-implantation changes of HFA and spike rates is shown in figure 2. In this example, HFA and spike rates declined over the initial ~100 days post-implantation before stabilizing (figure 2A). This post-implantation variability of HFA and spike rates was observed across most electrodes (figure 2B) and occurred in most patients (figure e-2). In addition, we also observed that the spatial distributions of HFA and spike rates changed during the initial days post-implantation (figure 2C). The spatial distributions of HFA and spike rate immediately following implantation do not necessarily reflect their spatial distributions after recovery from surgery (figure 2D).

**Figure 2.**
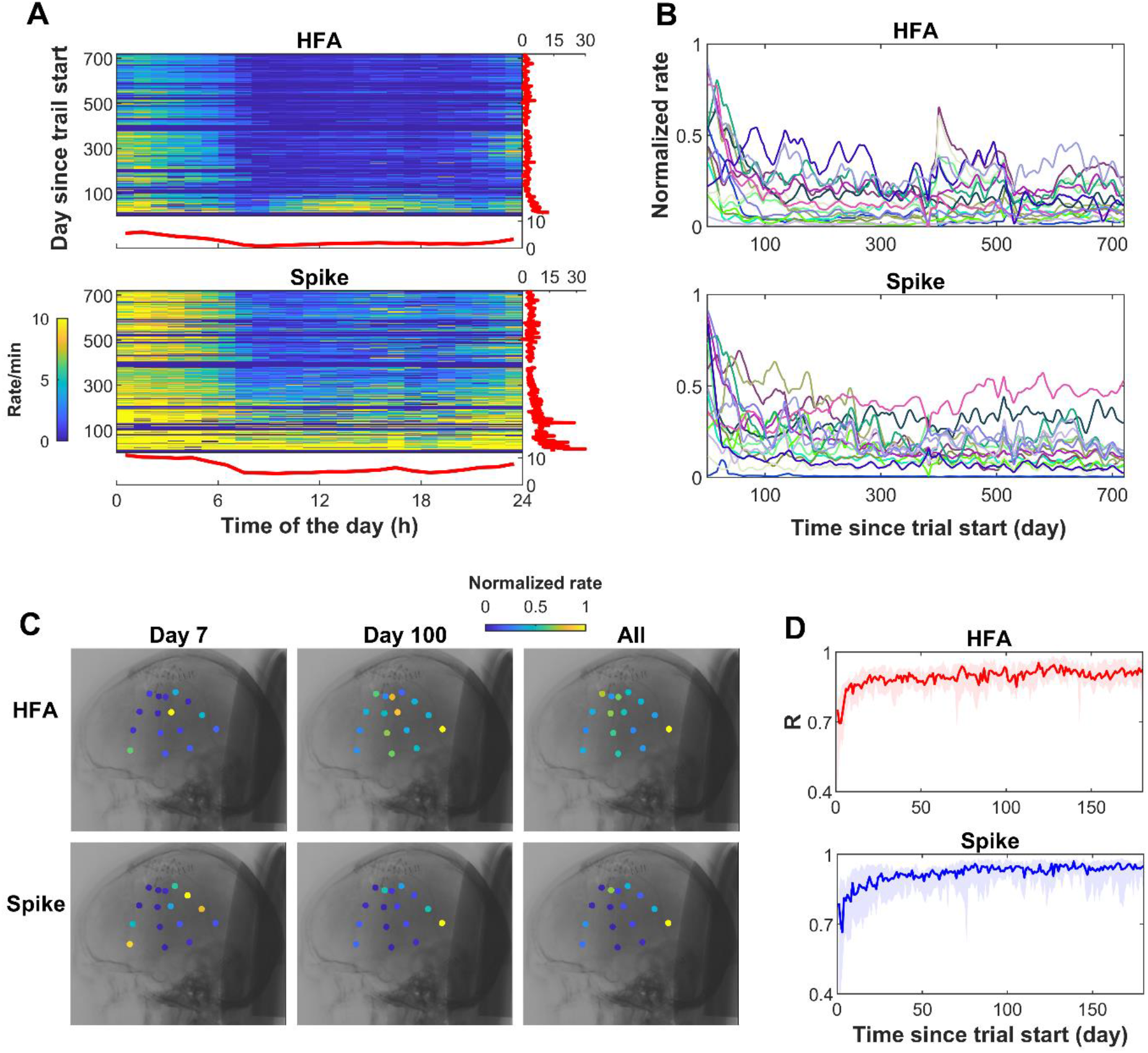
Post-implantation changes of rates and locations of HFA and spikes. (**A**) Example electrode of HFA and spike rate temporal distributions for each hour of a day since the beginning of the trial from one patient. The subplots to the right show the daily rates. The subplots below show the mean hourly rates averaged across all recording days. (**B**) Normalized HFA and spike daily rates across all recording days for each of the 16 recording electrodes from one patient. (**C**) Example of the post-implantation variability of HFA and spike rate spatial distributions from one patient. Rates were normalized to [0 1] based on their maximal values. HFA and spike daily rate spatial distributions at day 7 do not exactly represent their spatial distributions averaged across all recording days, while spatial distributions at day 100 are more similar to the spatial distributions averaged across all recording days. (**D**) *R* represents the Spearman correlation (i.e. similarity) between the spatial distributions of HFA/spike daily rate at each recording day and the spatial distributions of HFA/spike daily rate averaged across all recording days. The line indicates the median *R* value when summarized across all patients. The shaded areas represent corresponding interquartile (25-75%) ranges. The *R* value increases over the initial days post-implantation before stabilizing.

### Circadian and multiday cycles

ACFs were computed on the HFA and spike rates to determine if cyclic patterns existed. The ACFs over 48-hours are shown in figure e-3, and figure 3A illustrates these ACFs plots for four representative patients. All patients had prominent peaks in the ACF every 24 hours, demonstrating that both HFA and spike rates had strong circadian cycles. Some ACFs showed decaying trends, indicating they might have long-term trends or cycles with longer periods (e.g. Patient 10).

**Figure 3.**
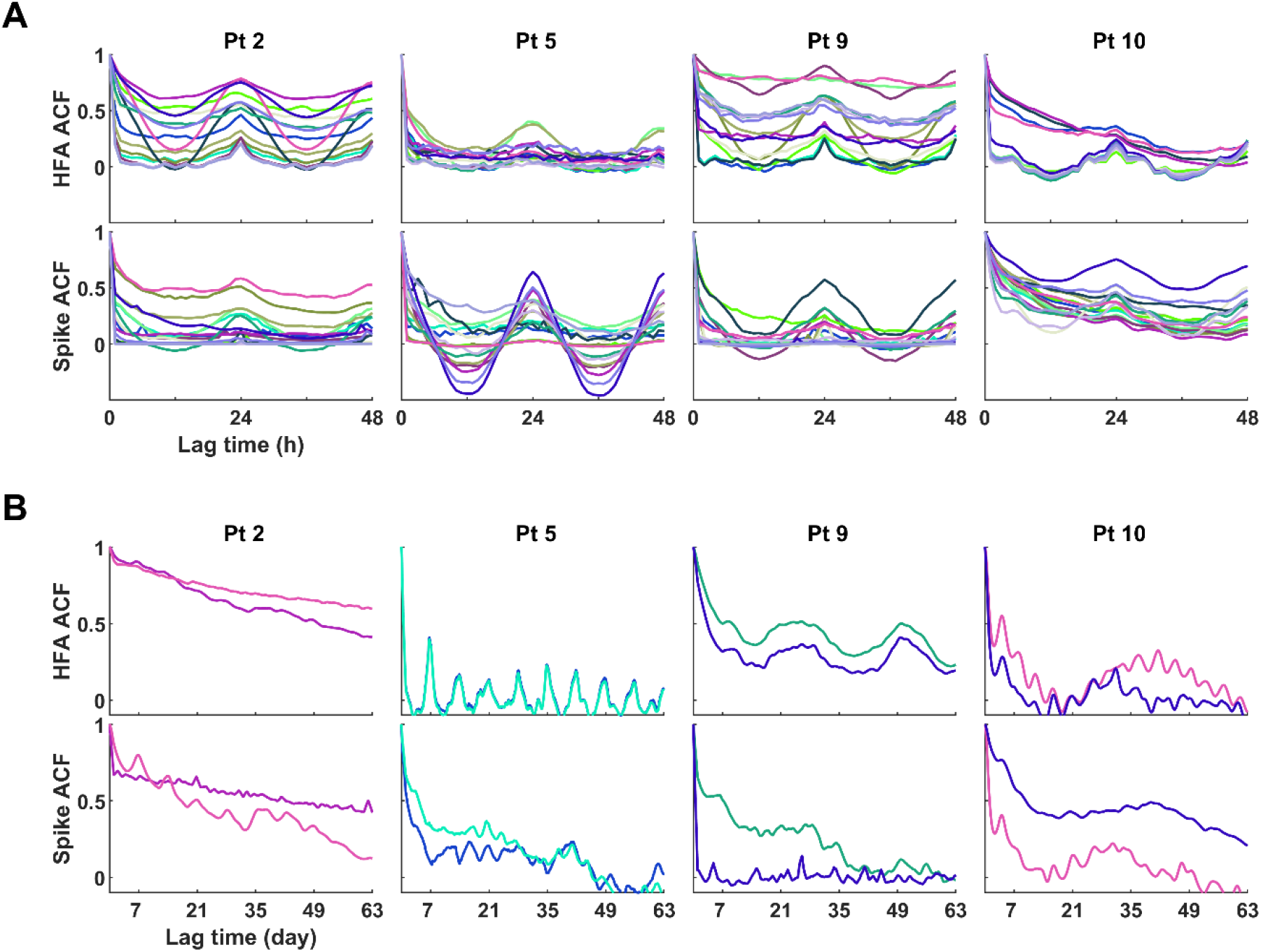
Examples of circadian and multiday cycles for four representative patients. (**A)** The autocorrelation functions (ACFs) of HFA and spike rates showing lag times up to 48 h. The different colors indicate each of the 16 electrodes; colors are consistent between HFA and spike plots. (**B)** The ACFs of HFA and spike rates on two representative electrodes showing lag times up to 63 days. Before doing the autocorrelation analyses, rate data were filtered using a 24 h moving average window to smooth out circadian fluctuations and highlight longer-term trends. The different colors indicate different electrodes; colors are consistent with plots in (A).

It has been previously reported that seizures are modulated by multiday cycles for some patients.^27^ We explored whether HFA and spike rates were also modulated by similar multiday rhythms. In general, the multiday cycles of HFA and spike rates were less obvious than circadian cycles. Only certain electrodes in some patients demonstrated multiday cycles. Figure 3B illustrates the multiday cycles in four representative patients. A weekly spike rate cycle was observed for Patient 2, and a strong weekly HFA cycle was observed for Patient 5. Longer cycles were also observed. A ~25-day HFA cycle was observed for Patient 9 and a monthly cycle was observed for Patient 10.

A summary of the detected circadian and multiday cycles across patients is shown in figure 4. These plots depict all the peaks derived from the ACF of the signal with significant cycles. All patients showed strong and robust circadian cycles for both HFA and spike rates. Twelve out of 15 patients showed multiday cycles for both HFA and spike rates, which tended to be limited to specific electrodes. Precise 7-day cycles were observed in some patients (e.g. HFA rates in Patient 5) while others had cycles of around 4-6 days (e.g. HFA rates in Patients 9, 10 and 11).

**Figure 4.**
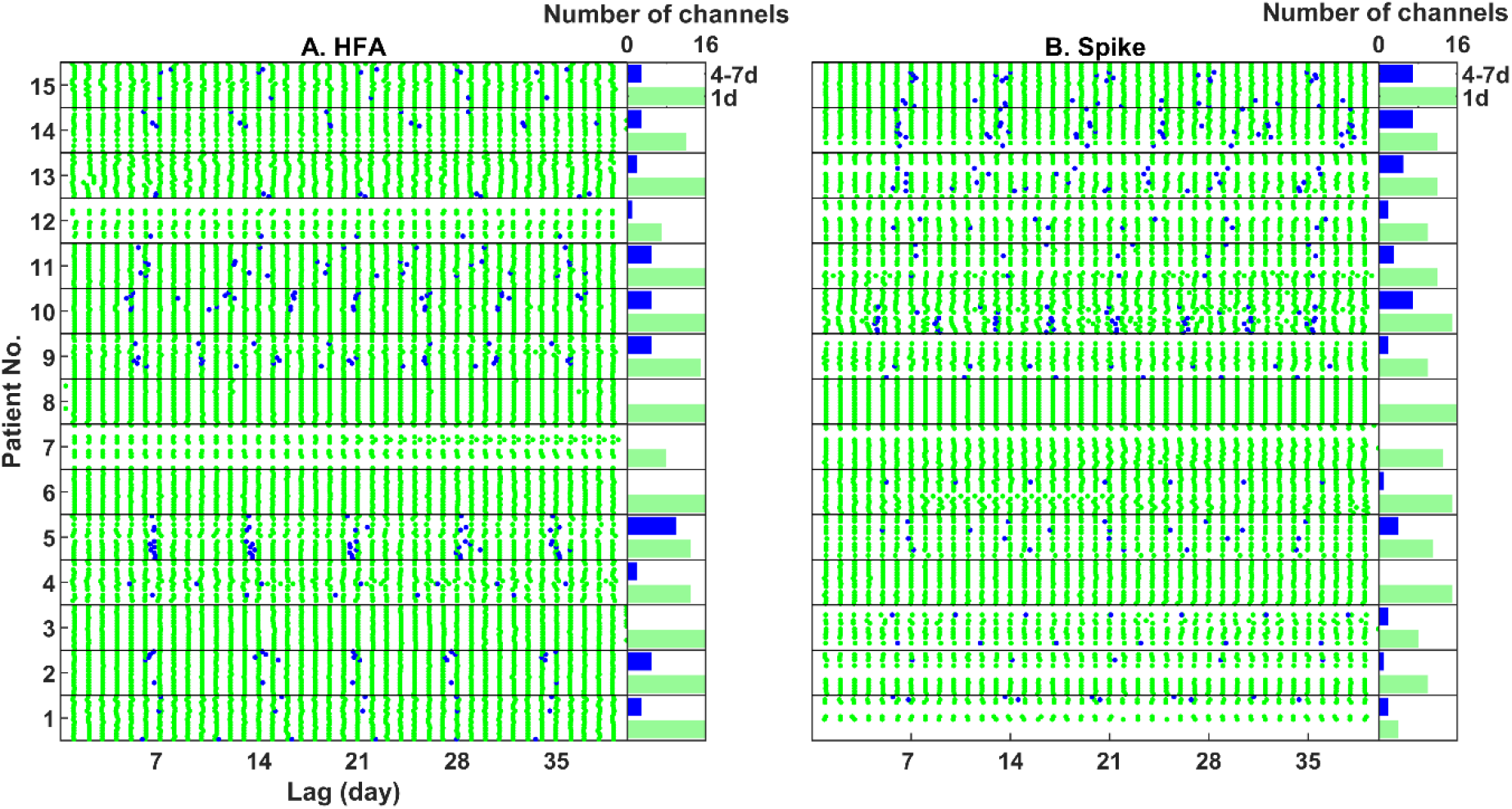
Summary of circadian and multiday cycles. For each patient, peaks and cycles were detected for each of 16 electrodes for (**A**) HFA rates and (**B**) spike rates. Note that, for each patient, there are 16 rows representing each of the 16 recording electrodes. The green dots represent the peaks of the ACF detected from the signal with significant short cycles, and the blue dots represent the peaks of the ACF detected from the signal with significant long cycles. The bar plots to the right of the HFA and spike plots represent the numbers of electrodes with significant cycles.

### Comparison of circadian patterns of HFA, spikes and seizures

The circadian fluctuations were patient-specific and often differed across electrodes. Figure 5 illustrates the circadian fluctuations of the HFA and spike rates in four representative patients.

**Figure 5.**
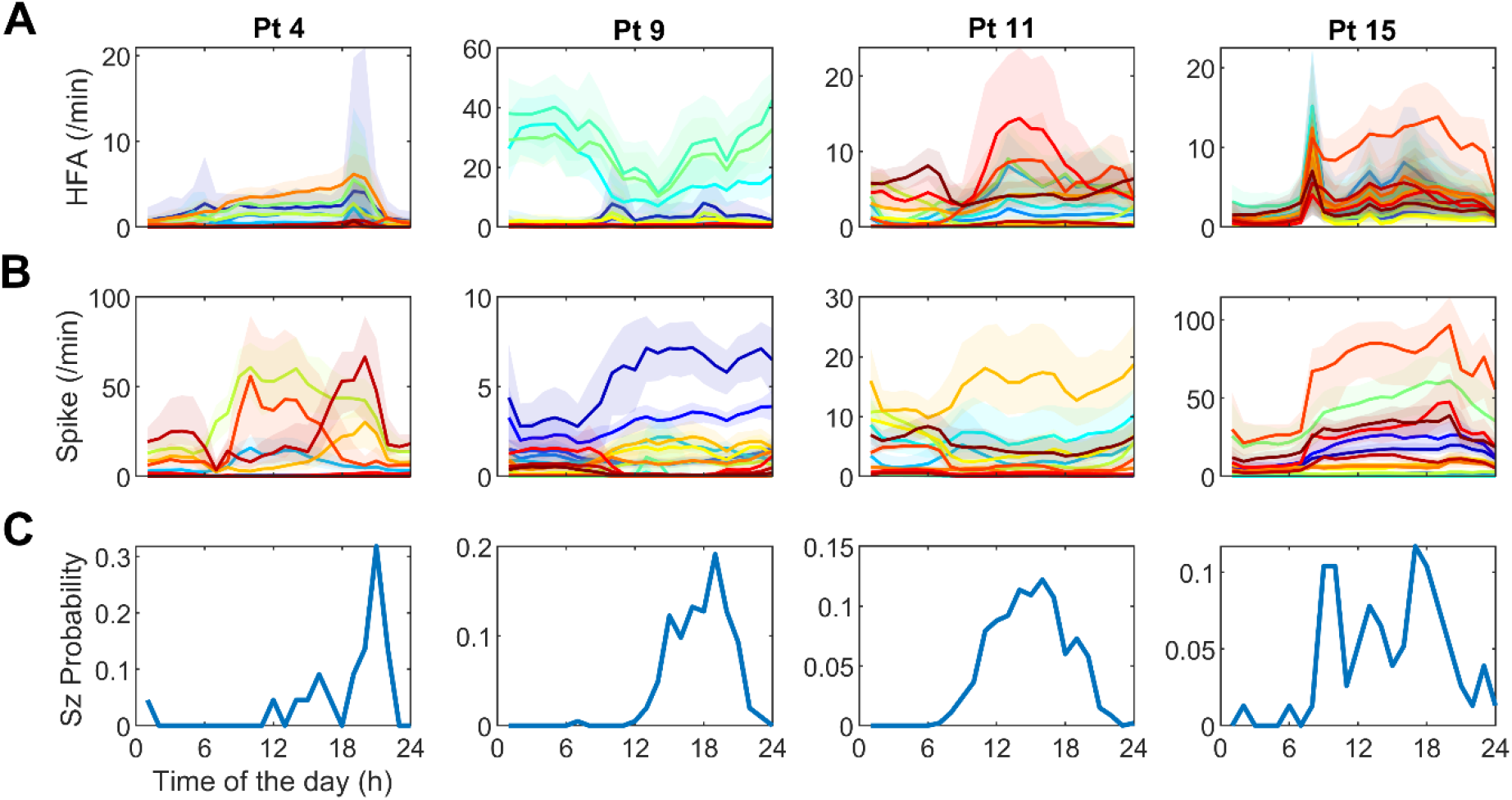
Circadian fluctuations for four representative patients. (**A**) HFA rates, (**B**) spike rates, and (**C**) seizure probabilities. Each colored line in (A) and (B) shows the median rate for each of the 16 electrodes; the shaded areas represent corresponding interquartile (25-75%) ranges. The different colors indicate different recording electrodes and are consistent between HFA and spike rate plots.

The circadian patterns of HFA rates often differed from spike rates, demonstrating that our detected HFA was physiologically different events from spikes. For instance, in Patient 9, the electrodes with high HFA rates were not the same electrodes with high spike rates, and they had out-of-phase circadian patterns. Furthermore, for some patients, circadian patterns varied across electrodes within individual patients (e.g. Patient 9), while in other patients, similar circadian patterns were observed across all electrodes (e.g. Patient 15).

When comparing the circadian patterns of HFA and spike rates with the seizure probability, we observed patient-specific behaviors (figure 5). In Patient 15, both the HFA and spike rates were phase-aligned with seizures in all electrodes. However, for Patient 11, HFA and spike rates were phase-aligned with seizures on some electrodes and anti-phase on others.

The circadian patterns of HFA and spike rates had statistically significant non-zero correlations with the circadian patterns of seizure probability in most patients (figure 6). There were statistically significant patient variability in correlation values between HFA and seizures, spikes and seizures (HFA: *F*=31.454, *p*<0.001; spike: *F*=5.405, *p*<0.001), and spike correlations had higher spatial variability than HFA correlations (spike: 78.41%; HFA: 34.44% of the total variability was caused by within-patient variability).

**Figure 6.**
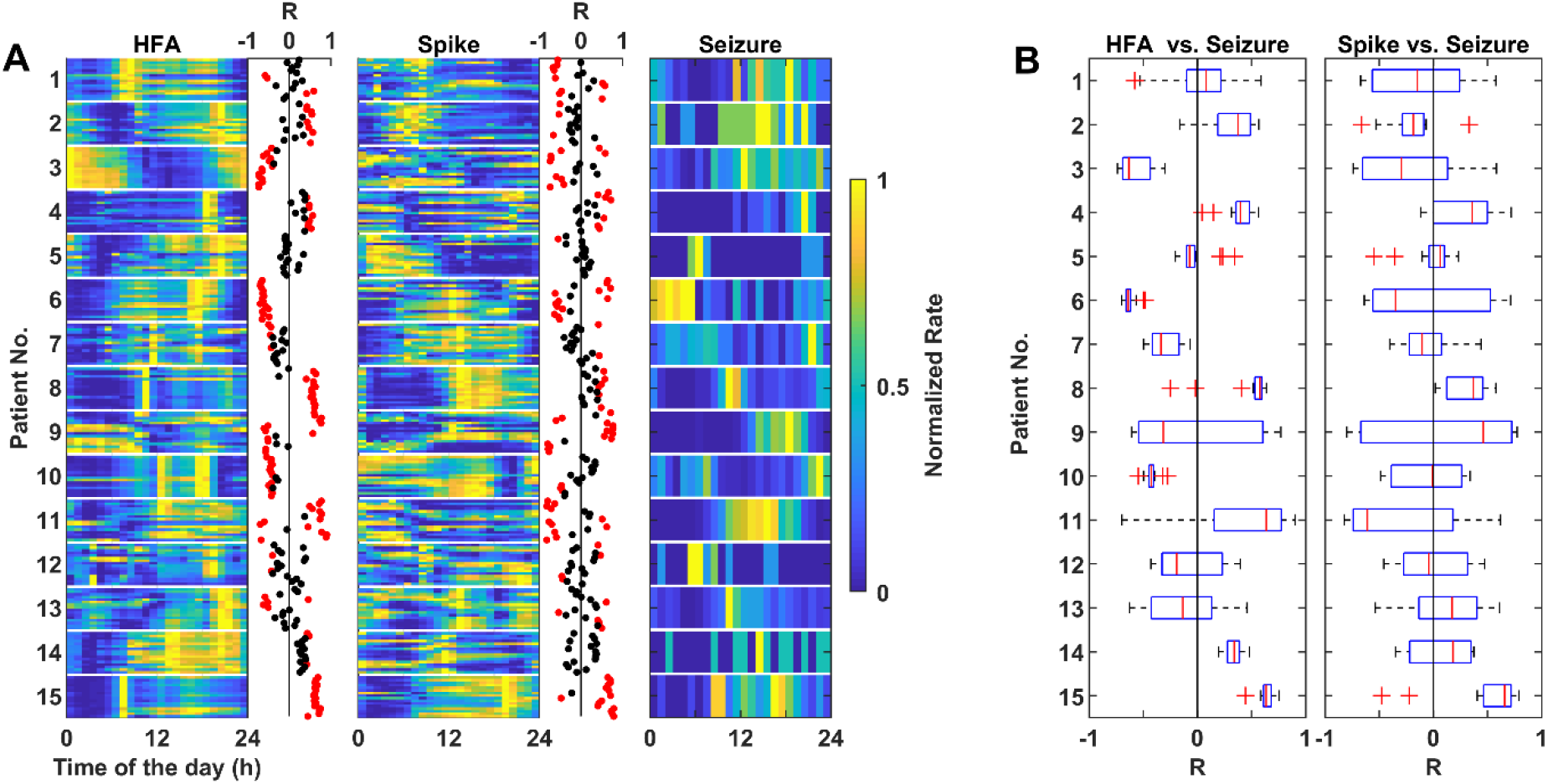
Comparison of circadian patterns of HFA, spikes and seizures. (**A**) The circadian fluctuations of the mean hourly rates of HFA, spike, and seizure averaged across all recording days. For each patient, there are 16 rows representing the 16 recording electrodes. For visualization purposes, all rates were normalized to the range [0 1]. *R* represents the correlation between mean hourly rates of HFA and seizures or the correlation between mean hourly rates of spikes and seizures. Red dots represent correlation values that are statistically different than zero (*P*<0.05), otherwise, they are shown as black dots. (**B**) Boxplots of the correlation values (*R*) for mean hourly rates of HFA vs. seizures and mean hourly rates of spikes vs. seizures for each patient. Values are presented as medians (the red middle line) with the interquartile range (ends of the boxes). Red + represent outliers.

### Validation

To validate that the observed rhythms were not influenced by changes during the first ~100 days following implantation, we removed the data from the first 100 days and recomputed the autocorrelations. The ACF remained similar (figure e-4), indicating that rhythms were robust despite the initial instability.

To validate that our HFA results were not driven by spikes, we removed any HFA that coincided with spikes and recomputed the autocorrelations. The results remained similar (figure e-5), indicating that the HFA circadian rhythms were independent of spikes.

Furthermore, the circadian rhythms were also independent of seizure states, as removing any HFA or spike that occurred during seizures produced similar ACF plots (figure e-6).

Finally, we validated that our results were robust to possible false detections or artefactual events by randomly deleting half of the HFA and recomputing the ACF. This process was repeated 1000 times to generate a distribution of HFA rates (figure e-7). The results indicated that the circadian cycles were robust to possible false detections in the automated detection process. These results were also robust to changes in the amplitude threshold used to detect HFA (figure e-8).

## Discussion

The HFA that we detected in this study had a relatively high similarity to the visually marked HFOs (table e-1). And this similarity value is comparable to the interrater agreement in HFO marking (table e-2). Therefore, while our detected HFA is more general than HFOs, our HFA encompasses most HFOs and generalizes well to HFOs marked by different reviewers.

This is the first study to explore the spatiotemporal patterns of HFA rates and compare them with spikes and seizures using long-term continuous iEEG recordings. We found a high amount of intra- and inter-patient variability in the rates of HFA and spikes. HFA and spike rates varied significantly throughout the day and varied significantly across electrodes. Furthermore, the spatiotemporal behavior of HFA and spikes immediately following electrode implantation did not reflect their long-term behaviors after recovery from surgery. These results caution against using HFO rates as a pre-surgical metric when only used at a specific hour of a day, and further caution about the effects of surgery itself. The variability resulting from post-implantation effects may account for the contradictory HFO results observed across many studies.^8, 9, 11, 12, 26^

Our findings of post-implantation changes of rates and locations of HFA and spikes agree with previous findings of post-implantation fluctuations of iEEG signals^13, 14^ and SOZ.^28^ These observations indicate that the fundamental mechanisms underlying epileptic networks seem to be altered after electrode implantation^13^ and one should be cautious when interpreting results using the EEG biomarkers extracted immediately following implantation. The cause of these changes may be related to the immunoreactivity after electrode implantation.^29^ There are seemly contradictory reports on whether HFO rates can be used to delineate the EZ.^5, 8, 12, 26^ Most existing HFO studies have used short-term recordings but may not be representative nor sufficient to present their long-term behaviors after recovery from surgery. Recently, Gliske et al. found that HFO locations were inconsistent over different hours or days of 10-min epochs of sleep iEEG in most of the patients.^8^ Fedele et al. found that HFOs could reliably predict surgical outcomes when HFO locations were consistent over time.^11^ Therefore, this study cautions against only relying on HFO or spike analysis for identifying the EZ.

Cyclic rhythms of similar durations in the rates of HFA, spikes, and seizures suggest that they may be co-modulated by factors operating at multiple timescales, such as antiepileptic drugs,^30^ sleep-wake cycles,^17, 25^ hormones,^31, 32^ stress,^33^ and even weather^34^. For instance, in epileptic rats, rates of HFOs, spikes, and seizures have been shown to decrease following the introduction of antiepileptic drugs.^35, 36^ In epileptic patients, the withdrawal of antiepileptic drugs caused an increase in HFO rates and seizures, while spike rates remained unchanged^37, 38^ or decreased.^39^ Both HFOs and spikes have been shown to occur more often during non-rapid eye movement sleep than during rapid eye movement sleep and wakefulness.^17, 25^ HFA in the seizure onset electrodes was shown to be strongly regulated by sleep-wake cycles, while HFA in non-seizure onset electrodes was not.^4^ It is also possible that some HFA may reflect physiological activities, such as memory consolidation.^40, 41^ Identifying the variables regulating HFA and spikes could provide feasible ways to investigate the underlying mechanisms that modulate seizures and may even suggest new management strategies by modifying those factors. Once the influence of confounding factors is better understood, they can also be taken into account by algorithms to increase their predictive performance.

Comparing the circadian patterns of HFA and spike rates to seizures based on the individual patient and individual electrodes extends our knowledge of their relationships. In contrast to previous reports,^22, 42, 43^ our study found a striking difference in circadian patterns of HFA and spike rates across electrodes within patients. This spatial variable behaviors of HFA and spike rates could help reconcile the discrepancies in the literature regarding how spikes and HFOs change prior to seizures.^2, 22, 38, 44–47^ In our study, we found statistically significant non-zero correlations between circadian patterns of HFA and seizures, and spikes and seizures in most patients (figure 6), indicating they are not entirely independent processes. However, the inter-patient and intra-patient (i.e., different electrode locations) variability in their correlations (figure 6), challenges the notion of a simple, generalizable relationship between EEG biomarkers and seizures. Future studies with long-term recordings and pathophysiology information of the individual electrodes may help better understand their relationships.

There are several limitations to our study. All the patients included in this study have drug-refractory focal epilepsy and, consequently, our findings may be limited to this type of epilepsy. Sleep, medications, pathophysiological information and other aforementioned variables may be useful to explain the patient-specific spatiotemporal patterns, but this is a complex question, which requires further investigation. However, with this unique long-term dataset, valuable insights have been derived which may also suggest future research directions. Also, our findings were limited to HFA. However, since there is no clear definition of HFOs that can enable the reliable identification of HFOs by different reviewers, we used a more general high-frequency event definition. Our HFA validation results indicate that our HFA were physiologically different from epileptiform spikes and that high confidence can be placed on our findings. In the future, it would be interesting to investigate the value of HFA for seizure forecasting, which is an important subject worthy of further detailed analysis.

## Glossary

EZ: epileptogenic zone
HFA: high-frequency activity
HFO: high-frequency oscillation
iEEG: intracranial EEG
SOZ: seizure onset zone

## Acknowledgments

ZC acknowledges the support of the China Scholarship Council (201709120011). MIM acknowledges support from the Melbourne Neuroscience Institute. We thank Associate Professor Graham Hepworth from the Melbourne Statistical Consulting Platform for his support with statistical analysis.

## Appendix: Authors

| Name | Location | Contribution |
| --- | --- | --- |
| Zhuying Chen, MM | The University of Melbourne, Australia | study design; data collection, analysis and interpretation; statistical analysis; drafting and revising manuscript |
| David B. Grayden, PhD | The University of Melbourne, Australia | study design; data interpretation; revising manuscript; study supervision |
| Anthony N. Burkitt, PhD | The University of Melbourne, Australia | study design; data interpretation; revising manuscript; study supervision |
| Udaya Seneviratne, PhD | St Vincent's Hospital, Melbourne, Australia | marking HFOs; revising manuscript |
| Wendyl J. D'Souza, PhD | St Vincent's Hospital, Melbourne, Australia | marking HFOs; revising manuscript |
| Chris French, PhD | Royal Melbourne Hospital, Australia | marking HFOs; revising manuscript |
| Philippa J. Karoly, PhD | The University of Melbourne, Australia | data analysis; revising manuscript |
| Katrina Dell, PhD | St Vincent's Hospital, Melbourne, Australia | data analysis; revising manuscript |
| Mark J. Cook, MD | St Vincent's Hospital, Melbourne, Australia | study design; data interpretation; revising manuscript; study supervision |
| Matias I. Maturana, PhD | The University of Melbourne, Australia | study design; data collection, analysis and interpretation; revising manuscript; study supervision |

