## Supplementary tables and figures for "Spatiotemporal patterns of high-frequency activity (80-170 Hz) in long-term intracranial EEG"

### HFA and HFOs cross-validation

Table e-1. HFA and HFO cross-validation using seven 16-channels 24-min epochs (2688-min of iEEG data in total) that were randomly selected from seven patients. One experienced neurologist (US) marked all epochs. Manually marked HFOs were compared to automatically detected HFA.

| Epoch No. | Patient | HFA | HFOs | Proportion of HFOs detected as HFA | Proportion of HFA not marked as HFOs | Similarity |
|---|---|---|---|---|---|---|
| 1 | 3 | 7,692 | 6,767 | 0.84 | 0.15 | 0.84 |
| 2 | 7 | 866 | 762 | 0.66 | 0.39 | 0.64 |
| 3 | 8 | 2,608 | 2,119 | 0.72 | 0.28 | 0.72 |
| 4 | 10 | 1,585 | 550 | 0.87 | 0.66 | 0.47 |
| 5 | 12 | 2,137 | 1,010 | 0.93 | 0.55 | 0.61 |
| 6 | 13 | 5,528 | 2,311 | 0.93 | 0.51 | 0.62 |
| 7 | 15 | 1,501 | 1,147 | 0.60 | 0.49 | 0.55 |
| Total | | 21,917 | 14,666 | 0.82 | 0.37 | 0.71 |

Table e-2. Independent HFO marking in the same 112 minutes of iEEG data by three reviewers (US, WD, and CF). Manually marked HFOs were compared to the automatically detected HFA and HFOs marked by different reviewers were also compared.

| Reviewer | HFA | HFOs | Proportion of HFOs detected as HFA | Proportion of HFA not marked as HFOs | Similarity / Inter-rater Agreement |
|---|---|---|---|---|---|
| US | 1,334 | 886 | 0.81 | 0.38 | 0.70 |
| WD | 1,334 | 738 | 0.82 | 0.51 | 0.61 |
| CF | 1,334 | 1,440 | 0.64 | 0.26 | 0.69 |
| US vs. WD | - | - | - | - | 0.75 |
| US vs. CF | - | - | - | - | 0.72 |
| WD vs. CF | - | - | - | - | 0.61 |

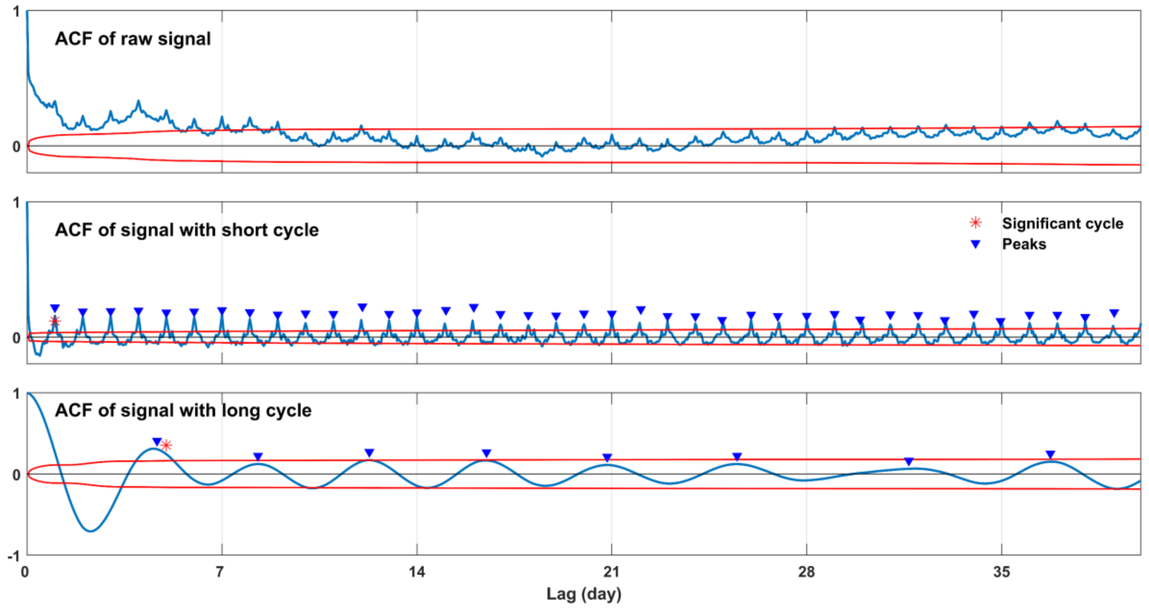


Figure e-1. Illustration of the multiple cycles detection method based on the regularity of peaks in the ACF of the filtered signal (raw signal is the HFA rate data from one example channel of Patient 10). The red curves show the 99% confidence bounds, an ACF value outside the bounds indicates that the ACF value is significant and is statistically unlikely to be caused by chance. * indicate significant cycles; triangles indicate peaks.

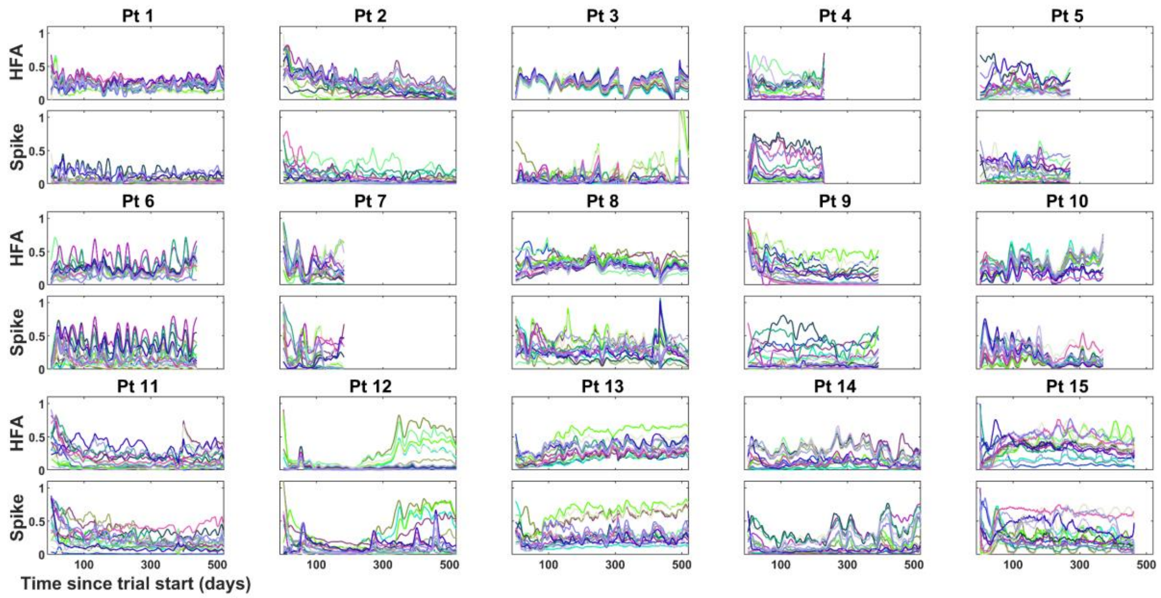


Figure e-2. Post-implantation variability of normalized HFA and spike daily rates for each patient and each of the 16 recording electrodes. For visualization purposes, normalized rates were smoothed using a moving average filter with a window length of 20 days.

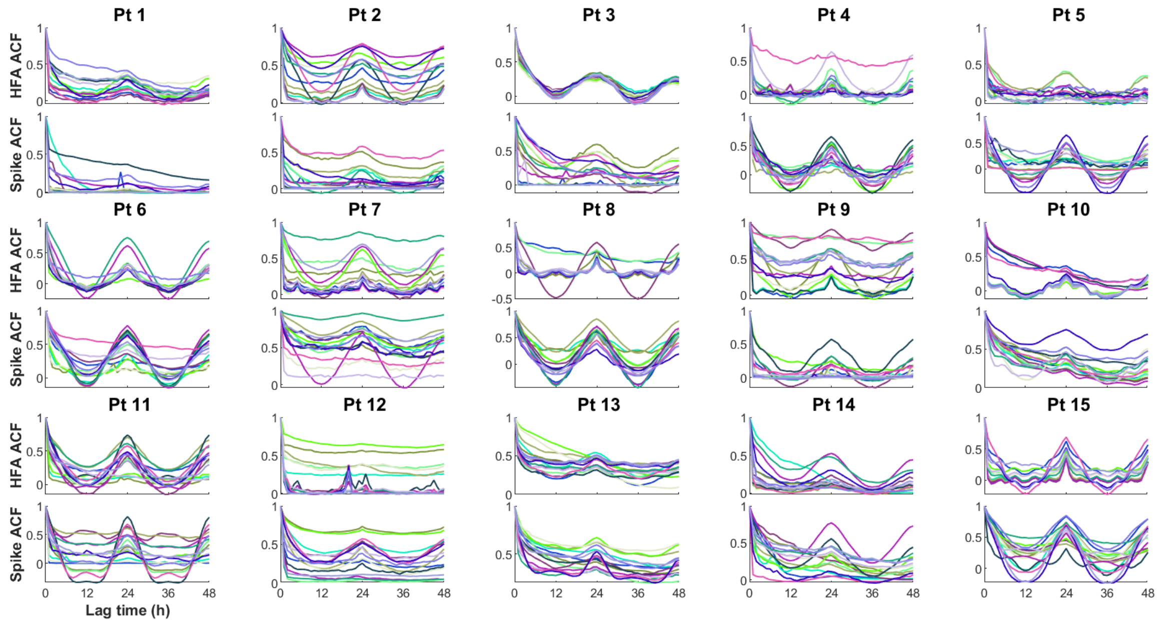


Figure e-3. The autocorrelations (ACFs) of HFA and spike rates showing lag times up to 48 h. The different colors indicate the 16 different electrodes; colors are consistent between HFA and spike plots.

### Validation

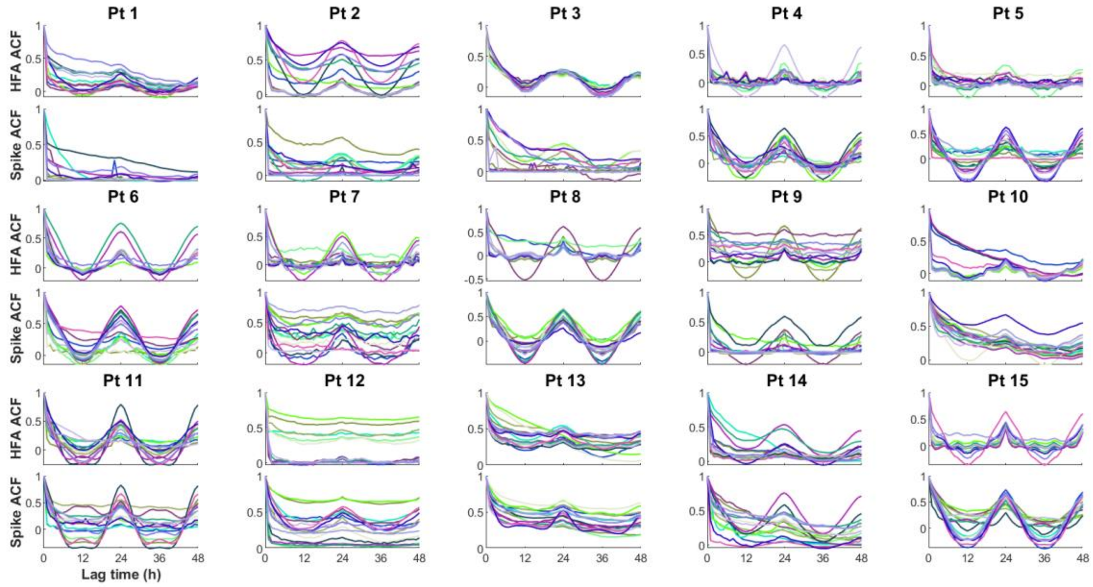


Figure e-4. The ACFs of HFA and spike rates (excluding data from the first 100 days) showing lag times up to 48 h. The different colors indicate different electrodes; colors are consistent between HFA and spike plots.

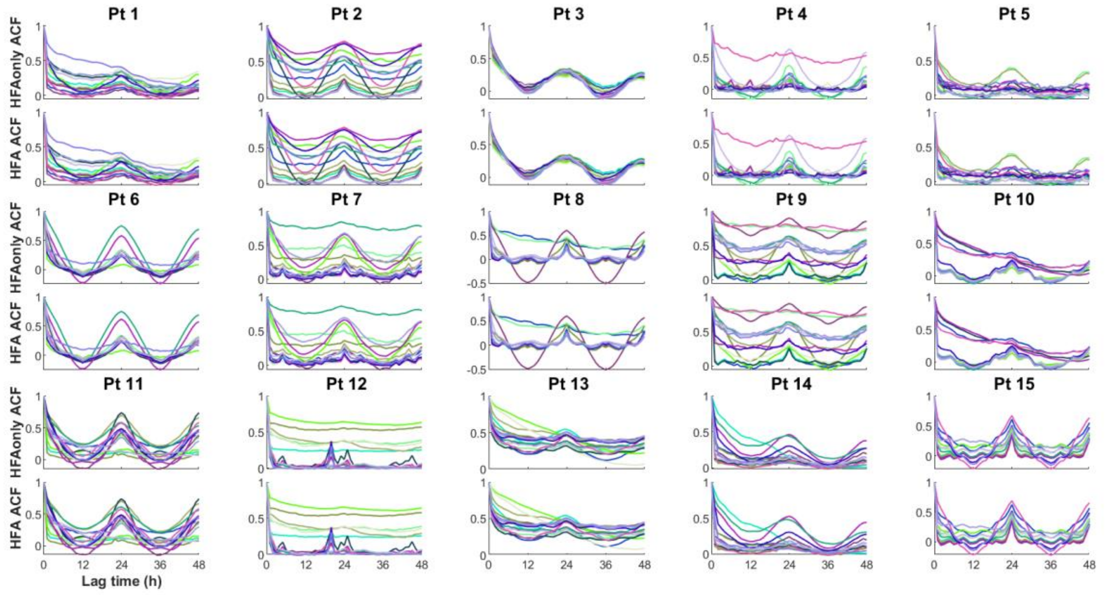


Figure e-5. The ACFs of rates of HFA without spikes (indicated as HFAonly) and HFA were compared. HFA without spikes were obtained by deleting any HFA indices that were up to 2 indices before the peak of spike and up to 8 indices after the peak of spike. The different colors indicate different electrodes; colors are consistent between HFA only and HFA plots.

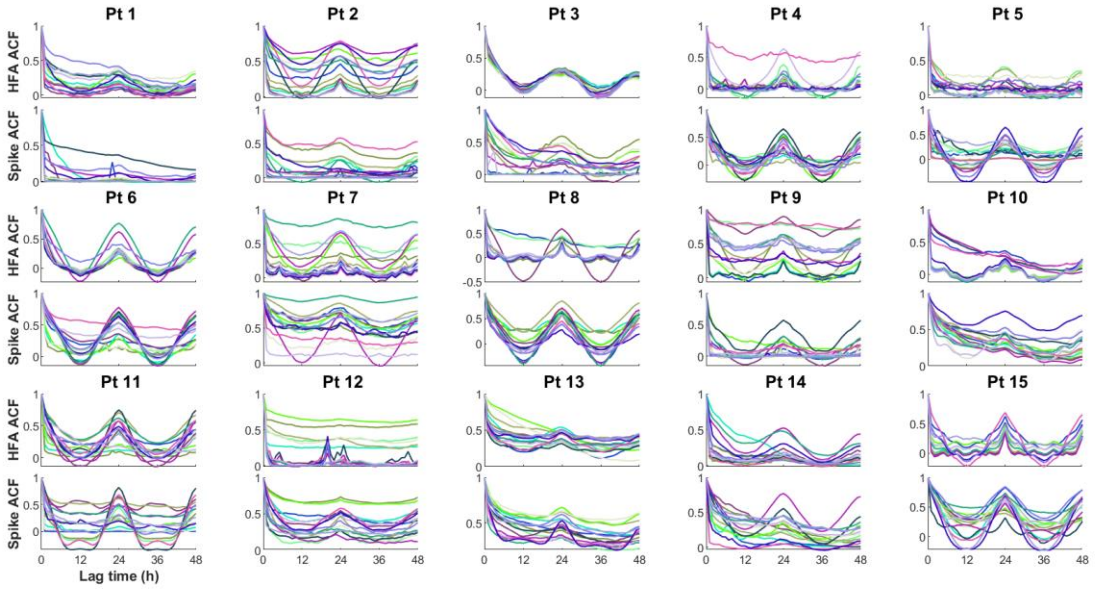


Figure e-6. The ACFs of rates of HFA and spike (excluding events in seizure periods) showing lag times up to 48 h. The different colors indicate different electrodes; colors are consistent between HFA and spike plots.

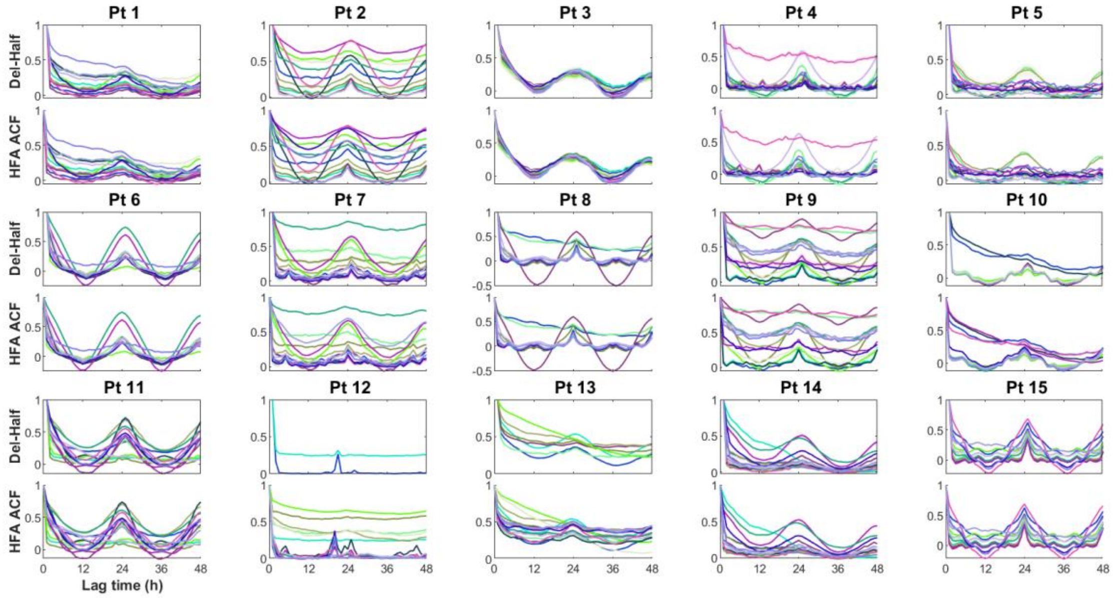


Figure e-7. The ACFs of HFA rates, where half of the HFA were randomly deleted and this process repeated 1000 times (indicated as Del-Half) compared to the original HFA rates. In the ACF plots of Del-Half-HFA rates, the different colored lines show the mean ACFs across the 1000 repeats; the shaded areas represent corresponding ranges (minimum, maximum).

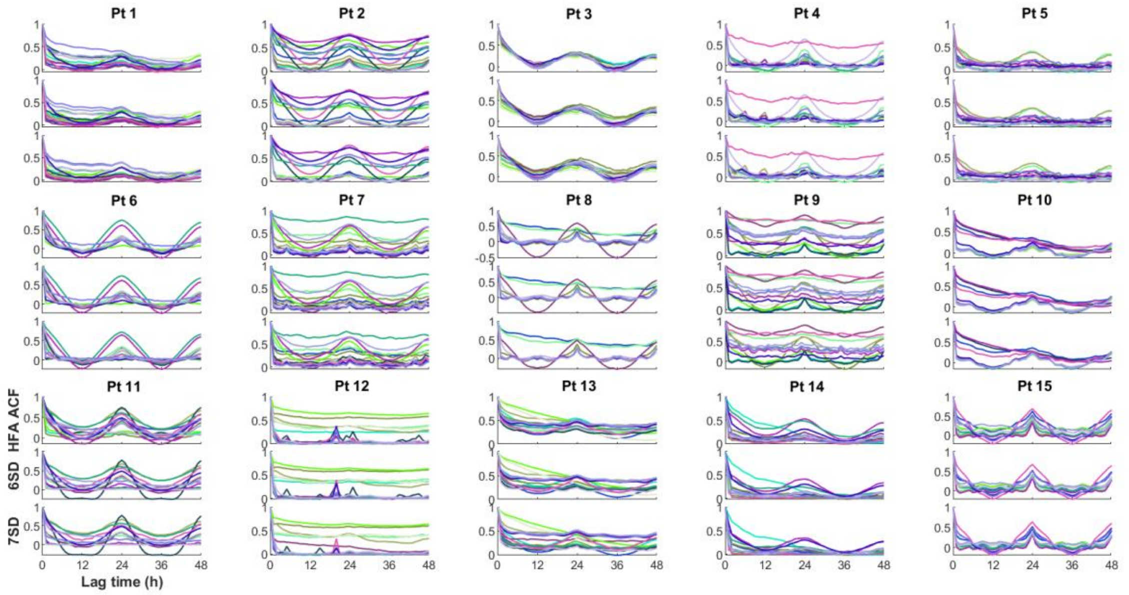


Figure e-8. Comparisons of the ACFs of HFA rates detected after increasing the amplitude thresholds to 6 standard deviations (SD) and 7 SD. The different colors indicate different electrodes; colors are consistent in plots of HFA detected by different thresholds.
